# A simple micropreparative gel electrophoresis technique for purification of proteins, nucleic acids, and bioconjugates

**DOI:** 10.1101/2021.03.26.436431

**Authors:** Sayyed Hashem Sajjadi, Shang-Jung Wu, Vitalijs Zubkovs, Hossein Ahmadzadeh, Elaheh K. Goharshadi, Ardemis A. Boghossian

## Abstract

The biochemical and biomedical fields hinge on the ability to effectively separate and purify biological macromolecules. Though this need is largely addressed with a variety of chromatographic and electrophoretic purification techniques, such techniques are usually laborious, time-consuming, and often require complex and costly instalments that are inaccessible to most laboratories. In this work, we introduce a simple micro-preparative (MP) method based on polyacrylamide gel electrophoresis (PAGE) to purify biological samples containing proteins, nucleic acids, and complex bioconjugates. Using a conventional vertical slab system, we demonstrate the extraction of purified DNA, proteins, and DNA-protein bioconjugates from their respective mixtures using MP-PAGE. We apply this system to recover DNA from a ladder mixture with yields of up to 90%, compared to the 58% yield obtained using specialized commercial devices. We also demonstrate the purification of folded enhanced yellow fluorescence protein (EYFP) from crude cell extract with 90% purity, comparable to purities achieved using a two-step size exclusion and immobilized metal-ion affinity chromatography purification procedure. Finally, we demonstrate the successful isolation of an EYFP-DNA bioconjugate sample that otherwise could not be processed using the two-step chromatography procedure. MP-PAGE thus offers a rapid and versatile means of purifying a variety of biomolecules without the need for specialized equipment.

## Introduction

The separation and purification of biological macromolecules are crucial not only for bioanalytics but also for advancements in biotechnology [1]. Macromolecules, in particular proteins and nucleic acids, can be isolated using a slew of techniques including high-speed centrifugation, membrane-based ultrafiltration, precipitation, electrophoresis, and chromatography [2–7]. These techniques differ in aspects such as scalability, throughput, yield, purity, precision, laboratory accessibility, and procedural complexity. Their advantages are used to identify the technique or combination of techniques that are most suitable for isolating a particular macromolecule under specific conditions for a given application. In fact, protein purification remains an evolving challenge that largely hinges on the researcher’s ability to piece together a suitable protocol based on existing techniques and optimization procedures. The diversity of these complementary techniques is therefore key to versatile protein isolation.

While flow chromatography systems especially benefit from relatively high throughput and scalability [5,8,9], several analytical applications such as sequencing and protein crystallography require only a small amount of purified protein. For such applications, techniques such as polyacrylamide gel electrophoresis (PAGE) are used to separate biomolecules according to their size and net electric charge [10,11]. The pore size in polyacrylamide gel (PAG) can be precisely controlled to separate biomolecules of a particular size with no need to use specialized equipment or columns for different size ranges, as performed in chromatography systems. In fact, PAGE serves as the gold standard for separating DNA with a single base pair resolution [12,13]. Due to its high resolution, low cost, and facile and rapid separation, PAGE is considered a standard and widespread analytical separation technique that can be found in most biological laboratories. In addition to DNA, this technique is especially useful for separating proteins either in their native or denatured forms [10].

While PAGE is typically used for analytical protein separation and identification, several studies have explored its use as a preparative procedure for purified protein isolation [14–23]. Large-scale (milligrams to grams of protein) electrophoresis can be achieved using preparative electrophoresis columns. However, the heat generated from this setup often leads to band broadening and protein denaturation [10]. For this reason, such setups require costly and specialized preparative gel electrophoresis equipment that is coupled with a cooling system. These separations are usually followed by post-elution methods for extracting the proteins from PAG after separation [24]. In addition, depending on the post-elution method, these extractions may require specialized setups, and the two-step separation and extraction procedure may alter protein activity and/or fold. Hao et al. [17] developed a novel microscale preparative electrophoresis system for protein separation. Although they confirmed high resolution and sufficient recovery of proteins based on gel electrophoresis, distinct separation and elution apparatuses was required. In some cases, specialized membranes are required for analysis. Fadouloglou [19], for instance, used a membrane in the electrophoresis setup to separate DNA. Although the custom-made configuration boasts advantages of its own, this customization requires a dedicated effort for constructing and troubleshooting the setup, which often limits the accessibility of the technology to specialized laboratories.

Despite the variety of preparative gel electrophoresis settings, previous techniques have been applied mainly to either DNA or proteins. Losses associated with complex and multi-step extractions often preclude the use of these techniques for the effective purification of dilute and low-yield samples, such as certain DNA-protein bioconjugates. While the Crush-and-Soak method represents the primary gel electrophoresis approach for bioconjugate purification [25], significant losses in the extraction step limit its use to concentrated samples. This method therefore prohibits the purification of a range of bioconjugate samples, including the dilute samples used in the study herein. Consequently, the purification of such dilute complexes has traditionally been limited to either chromatography techniques or multi-step approaches based on biomolecule tagging [26]. Zhou et al. [27] managed to purify DNA-protein conjugates through a complex, multi-step procedure requiring DNA modification and magnetic bead binding and release steps, resulting in the extraction of a modified conjugate containing a desthiobiotinylated tag. Yang et al. [28] had to rely on a sophisticated membrane-based approach for biomolecule isolation. Although this work presents a novel approach for separation, the large amount of impurities resulting from this method reveal ongoing challenges in biomolecule purification.

Herein, we present a single-step micropreparative (MP)–PAGE technique for double-stranded (ds) DNA and protein purification. The tri-layer slab gel arrangement, shown in Fig 1, utilizes a standard vertical electrophoresis setup available in most bioanalysis laboratories. The DNA extraction was compared to that achieved with the Crush-and-Soak method. In addition, the purities of the MP-PAGE protein samples were further analyzed and compared to those achieved with size exclusion chromatography (SEC) and immobilized metal ion affinity chromatography (IMAC) using recombinant enhanced yellow fluorescent protein (EYFP) as a model protein. Finally, the diminished losses and simplicity of this arrangement allowed us to apply MP-PAGE to a dilute and low-yield DNA-protein bioconjugate solution. This latter demonstration allows us to demonstrate, for the first time, a gel electrophoresis approach that can be used to purify dilute, low-yield bioconjugate samples that are otherwise intractable using existing gel electrophoresis techniques.

**Fig 1.**
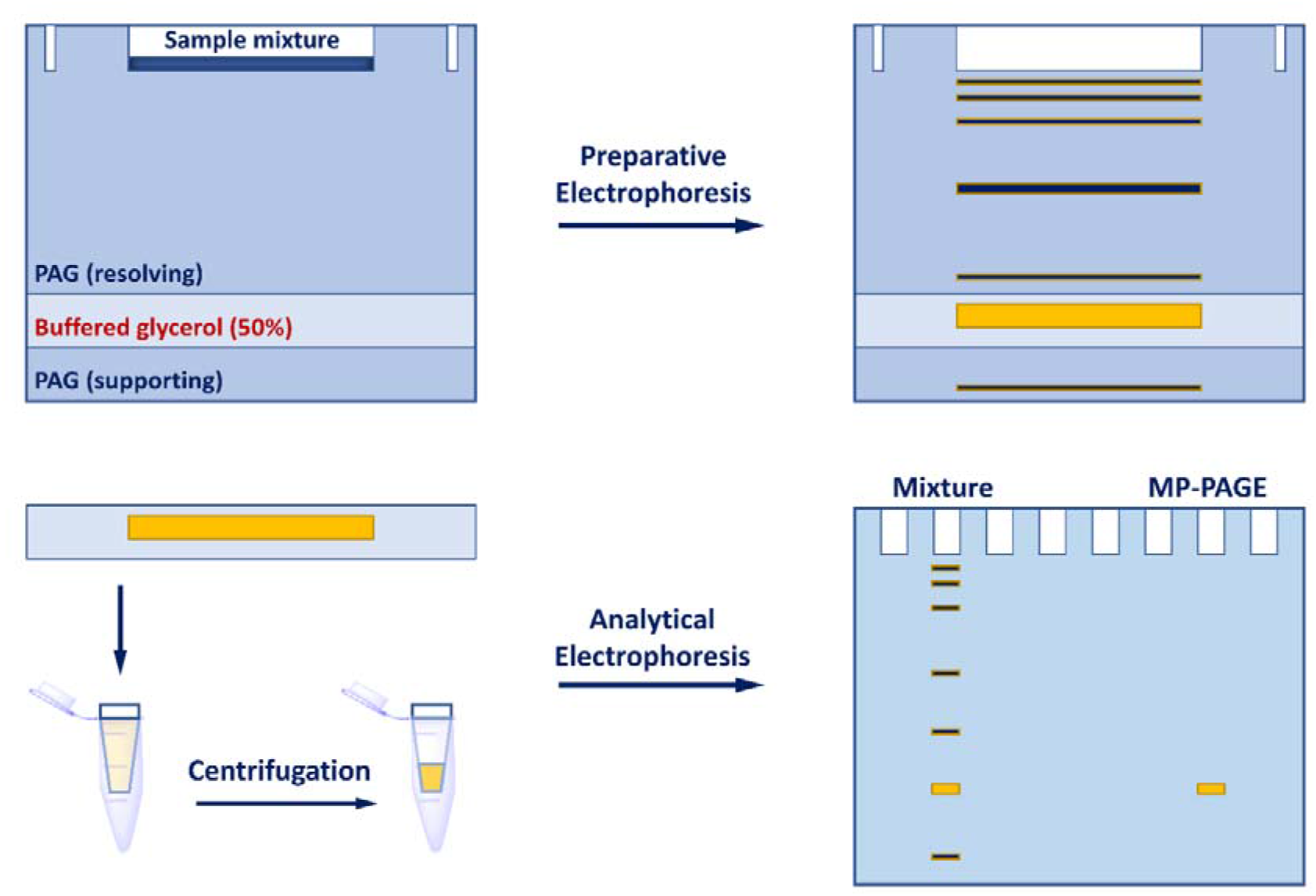
A schematic drawing of MP-PAGE for separating and extracting biological samples containing proteins and nucleic acids. The upper-left schematic shows the composition of a MP-PAGE gel containing a top resolving PAG, a glycerol layer for sample extraction, and a bottom supporting gel. The resolving gel is run at an adequate gel composition and voltage conditions for biomolecule separation. The band containing the desired biomolecule of interest (yellow band, top-right) is selectively eluted into the glycerol layer and run through a centrifugal filter (bottom-left) to remove the glycerol. An analytical gel is run on the extract (bottom-right) to confirm sample purity relative to the initial mixture.

## Materials and methods

### Preparation of EYFP enriched protein extract

Competent *E. coli* BL21 cells harboring the recombinant plasmid encoding EYFP were cultured in 2xYT medium supplemented with carbenicillin antibiotics. The cell culture was harvested and lysed for protein extraction. The insoluble materials were pelleted by centrifugation, and the supernatant (cell extract) was filtered through a 0.2 μm porous sterile filter and stored at 4 °C.

### EYFP purification using chromatography techniques: IMAC and SEC

IMAC purification was performed using a His-Trap HP 1 ml (GE Healthcare) on an ÄKTA START protein purification system (GE Healthcare). The sample was loaded in a buffer containing 20 mM phosphate buffer saline (PBS, pH 7.4), 500 mM NaCl, and 20 mM imidazole and then eluted in 20 mM PBS containing 500 mM NaCl, and 500 mM imidazole. 15.0 ml of the cell extract was loaded into the column and the chromatography was carried out at a flow rate of 1.0 ml/min for all steps (loading, washing, and His-tagged protein elution).

The SEC purification was achieved using a HiPrep 16/60 Sephacryl S-300 high resolution column (GE Healthcare) on the same ÄKTA system. The elution buffer contained 10 mM PBS (pH=7.4), and 140 mM NaCl. 3.0 ml of the sample was loaded into the column. The chromatography was carried out for 4 h with a flow rate of 0.5 ml/min. The sample mixture was separated and eluted into more than 40 fractions. The eluted fractions containing purified protein samples were concentrated using a 10 kDa Amicon centrifugal filter (Merck Millipore) and the elution buffer suspending the purified protein was replaced with EDTA (10 mM)-PBS (1x) buffer (pH=7.0) for long-term storage.

### EYFP conjugation to ssDNA

ssDNA-EYFP bioconjugation was performed as described previously [29]. The purified EYFP was reduced using 10 mM Tris(2-carboxyethyl)phosphine hydrochloride (TCEP, ABCR) in EDTA-PBS buffer for 1 h while shaking at 500 rpm in an Eppendorf thermomixer at 5 °C. The product was purified using a PD midiTrap G-25 desalting column (GE Healthcare) eluted with the same buffer. Freshly prepared 20 mM dibenzocyclooctyne-maleimide (DBCO, Tokyo Chemical Industry) in dimethyl sulfoxide was mixed with reduced EYFP (0.05 mM). The sample was shaken overnight at 5 °C, and the non-reacted DBCO was subsequently removed using a PD midiTrap G-25 desalting column. EYFP-DBCO was mixed with 5’-N_3_-GCA TGA ACT AAC GGA TCC CCT ATC AGG ACG-3’ (BH-N_3_, Microsynth) in a 1:4 molar ratio. The sample was left to react at room temperature accompanied by shaking for 4 h. A mixture of EYFP (without DBCO) and BH (without azide group) was incubated in the same way as the control.

### MP-PAGE separation of DNA and protein samples

The concentration of the PAG could be varied between 4 and 20%, depending on the size of the biomolecule of interest. The gel contained a 50% glycerol layer in running buffer for collecting the target band, followed by a supporting gel layer in running buffer (Fig 1). After sample loading, the electrophoresis was run until the band of the target molecule reached the glycerol layer. Prior to its elution into the glycerol layer, this layer and the supporting gel layer were replaced with new layers to remove undesired species with higher mobilities than the biomolecule of interest. The electrophoresis was re-applied until the band of interest was eluted into a new glycerol layer. The eluted sample was then recovered with a syringe and further concentrated using centrifugal filters for protein and protein-DNA conjugate samples and ethanol precipitation for DNA sample.

An acrylamide/bisacrylamide stock solution of 40% (19:1) was used to prepare PAGs in both preparative and analytical DNA-PAGE experiments while a 30% (37.5:1) solution was used for the protein-PAGE experiments. Both stock solutions were purchased from Carl Roth.

### DNA separation

A ds-DNA ladder (HyperLadder 25-500 bp, Bioline) was pre-stained with Sybr-Gold (SG, Invitrogen) and incubated in sample buffer (50% glycerol with 2x tris-boric acid-EDTA, TBE) for 30 min before injection into the gel. The pre-staining allowed us to track the DNA bands during electrophoresis. The preparative separation of DNA was performed on a MP-PAGE gel consisting of a 3 cm 12% resolving PAG and 2 cm 50% glycerol layer with 2x TBE buffer. After extraction of the glycerol layer, the separated DNA sample was concentrated by ethanol precipitation, which also removed the tracking dye. For comparison, the same bands were also separated using a conventional post-elution extraction procedure known as Crush-and-Soak [30]. For this procedure, the DNA mixture was run in a normal native PAG with the same concentration as the resolving gel in the MP-PAGE setup. The bands of interest were cut after electrophoresis using a scalpel, excised with a needle into small pieces, and soaked in buffer (1x TBE). The sample was incubated overnight, and the DNA that diffused out of the gel was concentrated by ethanol precipitation. The purity and recovery yield of the separated samples were compared using analytical electrophoresis on a 12% resolving PAG. To estimate the recovery yield, the corresponding volumes of the concentrated sample (taking into account the dilution factor for the recovered sample during ethanol precipitation) and original mixture were run on an analytical gel. The gel profiles were digitized using Image J software. The area of each peak was integrated and divided by the area of the corresponding peaks from the original mixture.

### Protein separation

MP-PAGE was used to purify EYFP from the crude cell extract. The preparative separation of EYFP was performed on a gel consisting of a 3 cm 12% native resolving PAG and 2 cm 50% glycerol layer with tris-glycine buffer. 300 μl of crude extract, containing ~1.6 mg of EYFP (according to its characteristic absorption peak at 514 nm with an extinction coefficient, *ε*_*EYFP*,514_, of 83400 M^−1^ cm^−1^ or 2.98 ml mg^−1^ cm^−1^ [31]), was loaded onto the MP-PAGE gel. The sample was run at 150 V until the fluorescent band migrated into the glycerol layer.

The purity of the protein samples extracted either via MP-PAGE or chromatography was compared using reducing sodium dodecyl sulphate (SDS)-PAGE in tris-glycine-SDS (TGS) buffer. 15 μl of each sample containing 1.0 μg of EYFP (based on *A*_514 nm_) in reducing sample buffer (containing dithiothreitol) was loaded onto a gel consisting of 4% stacking and 12% resolving PAGs. After 1 h of electrophoresis at 150 V, the gel was stained by Coomassie brilliant blue (CB).

The protein concentration of the extracted samples was determined from UV-visible spectra recorded using a NanoDrop 2000 (Thermo Scientific). 1.3 μl of each sample was loaded onto the device, and the spectrum was recorded in the 250 – 600 nm range. Total protein concentration was determined using a bicinchoninic acid (BCA) assay from the Pierce™ BCA Protein Assay Kit (Thermo Scientific). The fluorescence spectra of the EYFP samples were measured with a Varioskan LUX Multimode Microplate Reader (Thermo Scientific).

### DNA-Protein conjugate purification

The conjugate sample was purified using the same gel as described in the previous section (Protein separation). To assess the purity of the MP-PAGE conjugate, we performed reducing SDS-PAGE for 1 h at 150 V in TGS buffer (followed by CB staining) and urea-PAGE for 45 min at 200 V in TBE buffer (followed by SG staining) for protein and DNA detection, respectively.

## Results and discussion

### Separation of ds-DNA

We applied the MP-PAGE setup shown in Fig 1 to separate and extract individual dsDNA bands from a ladder mixture. In this setup, the desired bands were eluted and collected from the glycerol layer. Compared to buffered aqueous solutions used in specialized electrophoresis flow systems [19,24], the higher viscosity of the glycerol layer sufficiently impedes band mobility, allowing distinct DNA bands to remain separated in the glycerol layer. In addition, the denser glycerol layer shows limited diffusion into the PAG pre-solution, allowing the pre-solution to solidify in the presence of the neighboring glycerol layer.

This arrangement was used to separate a 25-bp band from a 12-strand ds-DNA mixture (Fig 2A). As shown in Fig 2A, we were also able to sufficiently separate 50 and 75-bp DNA strands using the same gel composition and operating voltage conditions, demonstrating the versatility of this technique in separating different DNA sizes from within a mixture. The calculated recovery yields were 90, 80, and 77 % for the 25, 50, and 75-bp DNA strands, respectively (Fig 2B). The gradual decrease in the yields with increasing DNA length is attributed to the resolving gel conditions optimized to favor separation of the 25 bp strand. The same DNA strands were also separated using the traditional Crush-and-Soak method with calculated recovery yields of 58, 54, and 24 % for the 25, 50, and 75-bp DNA bands, respectively. Compared to MP-PAGE, the Crush-and-Soak method showed lower extraction yields and an even greater decrease in the extraction of larger DNA strands under the tested conditions. This difference in DNA extraction is largely attributed to the diffusion-limited separation of the DNA from the solid gel matrix in the Crush-and-Soak extraction [32].

**Fig 2.**
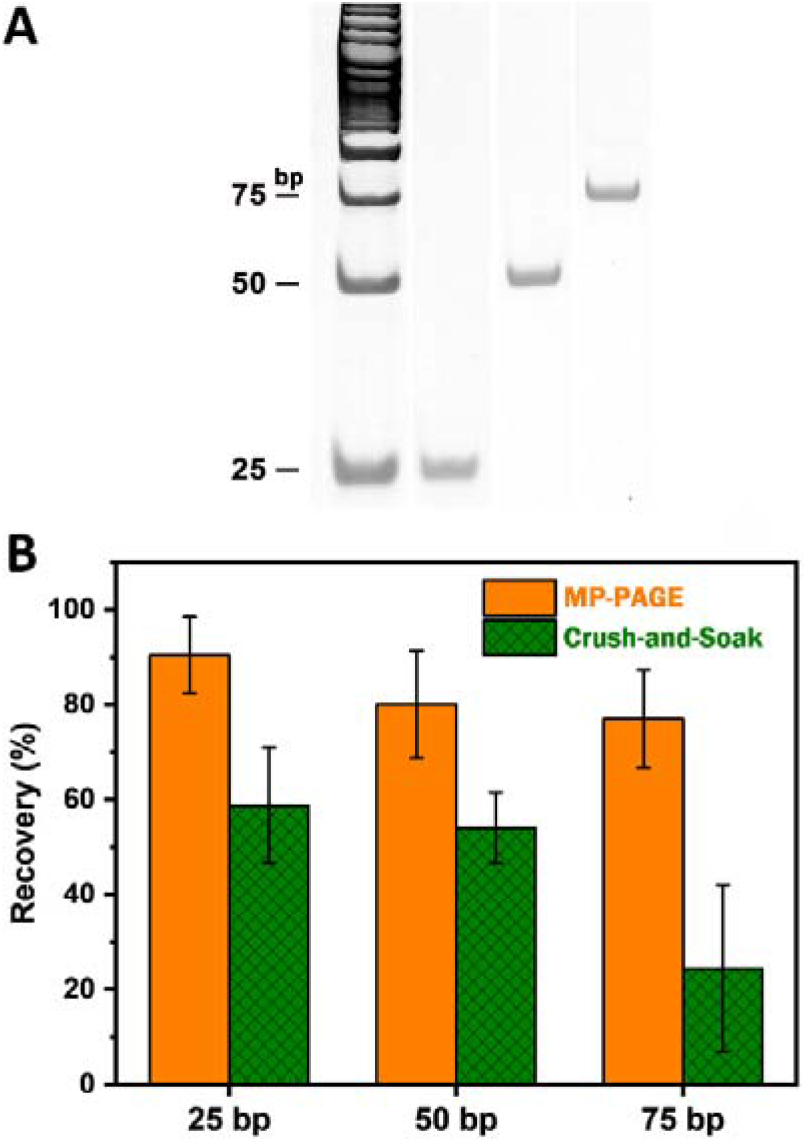
Separation of ds-DNA lengths from a ladder mixture containing 12 different DNA lengths. (A) 12% Native PAGE profiles (stained with SG) of the MP-PAGE-separated DNA samples. (B) Comparison of the purified DNA recovery yields from the MP-PAGE and the Crush-and-Soak techniques. Experiments were performed in triplicates, and the error bars represent standard deviations.

### Protein purification: separation of EYFP from E. coli extract

The MP-PAGE was also used to recover over-expressed EYFP from recombinant *E. coli.* Fig 3A shows the analytical reducing SDS-PAGE gel of the crude bacterial extract enriched with the recombinant EYFP, accompanied by a number of contaminating proteins ranging between 10 and 100 kDa. The EYFP is represented by a double band at around the expected size of 27 kDa. This double band is attributed to the cleavage of a signaling sequence for some of the proteins [33]. The proteins contained by the bands were extracted and purified using MP-PAGE, SEC, IMAC, as well as a double-extraction method based on IMAC extraction followed by SEC (chromatograms and corresponding CB stained gels shown in Figs. S1 and S2). Compared to the MP-PAGE sample, which only showed sparse, faint contaminating protein bands, the SEC sample contained a greater amount of both smaller and larger protein impurities under the tested conditions. The impurities were particularly pronounced for sizes neighboring the desired ~27 kDa band. In contrast, the IMAC extraction yielded a significantly reduced level of impurities with a pronounced contaminating protein band at around 10 kDa. These impurities are attributed to non-specific binding of histidine- or arginine-rich proteins that can intrinsically bind to the nickel column. Compared to SEC and IMAC alone, the double-extraction IMAC+SEC method yielded the least amount of contamination, showing only faint impurity protein bands. Based on the gel analysis, the sample purity was comparable to that achieved with MP-PAGE, except for a faint impurity band at 40 kDa that was more pronounced in the MP-PAGE sample. As such, the samples purified from the IMAC+SEC method were used in the subsequent analysis to compare the integrity of the proteins isolated with MP-PAGE.

**Fig 3.**
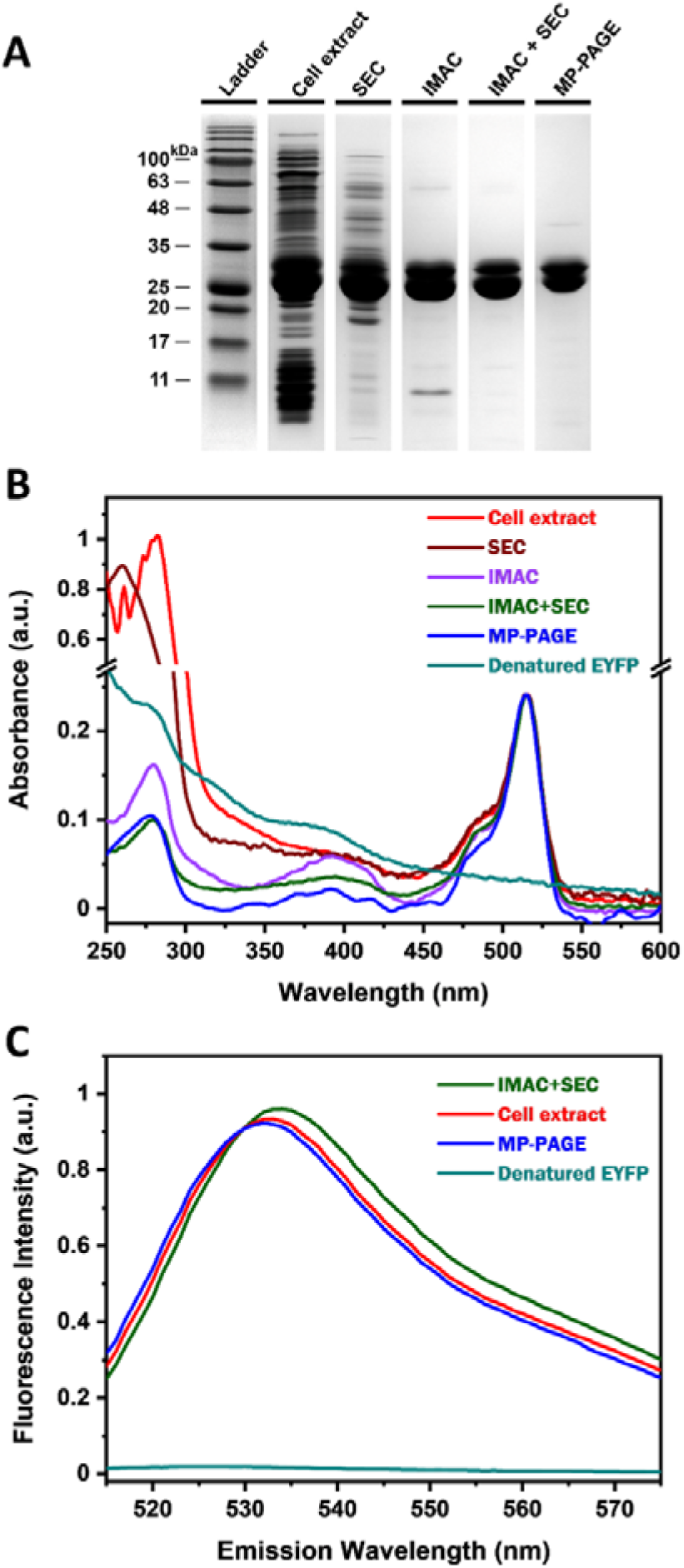
Analysis of protein-containing samples using gel electrophoresis, absorbance, and fluorescence spectroscopies. (A) CB-stained reducing SDS-PAGE profiles of crude cell extract and purified EYFP samples from SEC, IMAC, IMAC + SEC, and MP-PAGE. To compare the amount of impurities for an equivalent amount of EYFP, each lane was loaded with sample volumes that have the same EYFP peak absorption at 514 nm (*A*_514nm_ = 0.2). (B) UV-visible spectra of crude cell extract, purified EYFP samples achieved by SEC, IMAC, IMAC + SEC, and MP-PAGE, as well as denatured EYFP. Denatured EYFP corresponds to the IMAC+SEC sample heated to 95 °C for 5 min. (C) Fluorescence spectra of crude cell extract and purified EYFP samples from IMAC+SEC (as the purest protein sample) and MP-PAGE, as well as denatured EYFP described in (B). The samples were excited at 490 nm and fluorescence spectra were normalized by the EYFP concentration.

The purity and integrity of the samples were further analyzed using absorption and fluorescence spectroscopies. All extracted samples, except the desaturated EYFP, showed an absorption peak at 514 nm (Fig 3B). Furthermore, Fig 3C shows the fluorescence of EYFP with its characteristic peak at around 533 nm upon 490 nm excitation. Both absorption and fluorescence peaks were diminished when the samples were heated to 95 °C for 5 min. Since this treatment denatures EYFP, the diminished absorption and fluorescence peaks verify that the detected optical signatures correspond to properly folded protein. The absorption spectra were also used to calculate sample purity. Assuming a relatively pure protein solution from the IMAC-SEC sample, we calculated an effective EYFP extinction coefficient of *ε*_*EYFP*,280_ ≈ 1.23 ml mg^−1^ cm^−1^ based on total protein concentration (as determined by the BCA assay) of the IMAC-SEC sample and the optical absorption (*A*) measured at 280 nm, which corresponds to the non-specific absorption peak of proteins containing aromatic residues. Total native EYFP protein concentration was calculated based on its specific absorption peak at 514 nm. By comparing the calculated protein concentration at 514 nm with the effective concentration at 280 nm, the native-EYFP purity (*%P*) in the different samples was calculated using the following equation:

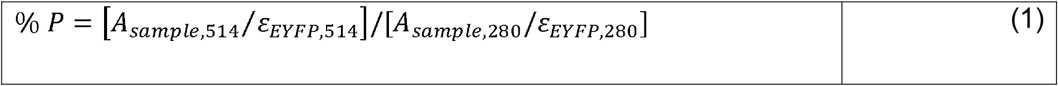

Based on this calculation, the protein purity was found to be 16, 58, 94, and 90% for SEC, IMAC, IMAC+SEC, and MP-PAGE, respectively.

### Purification of DNA-protein conjugate

DNA–protein conjugates are used in a variety of applications ranging from DNA-based nanofabrication to target recognition and signal amplification [27,34]. Several approaches have been developed to conjugate proteins and DNA [35]. However, the conjugation product often requires extensive treatment to remove unreacted precursors. Methods such as the Crush-and-Soak method, which was used for the DNA recovery described above, result in poor recovery yields [25]. Alternative methods such as the biotin displacement strategy developed by Zhou et al. [27], ion-exchange chromatography [36], and SEC [37] can achieve higher yields, though the extracted conjugates are often accompanied by sample impurities such as unreacted reagents as well as the need for specialized equipment and columns.

In our study, we attempted to purify a ssDNA-EYFP bioconjugate synthesized *via* azide-DBCO click chemistry. Specifically, ssDNA-azide was mixed with DBCO-EYFP to form a covalently linked ssDNA-EYFP based on stable triazole formation. The limited amount of sample could not be processed using SEC due to sample loss from significant dilution and the inability of SEC to discern unreacted proteins from their conjugates. However, the limited sample amount was sufficient for MP-PAGE purification. The successful conjugation of the ssDNA-EYFP was confirmed with urea-PAGE, which verified the formation of a new conjugate band following the reaction (Fig 4A). The conjugated sample showed decreased electrophoretic mobility compared to the non-conjugated EYFP due to the appended negatively charged oligonucleotide [38]. Conjugation was also confirmed with SDS-PAGE (Fig 4B) with a ~9 kDa increase in molecular weight corresponding to the attachment of one oligonucleotide per EYFP. Since the electrophoretic mobility of DNA (Fig 4A) and protein (Fig 4B) did not change in the absence of azide and DBCO, the new bands appearing in the samples originate from the covalent conjugation of EYFP-DNA. The presence of unreacted DNA and protein is confirmed by the SDS-PAGE even after filtration through a 30 kDa centrifugal membrane. In contrast, bioconjugated sample purified through MP-PAGE shows complete isolation of the conjugate from the unreacted compounds.

**Fig 4.**
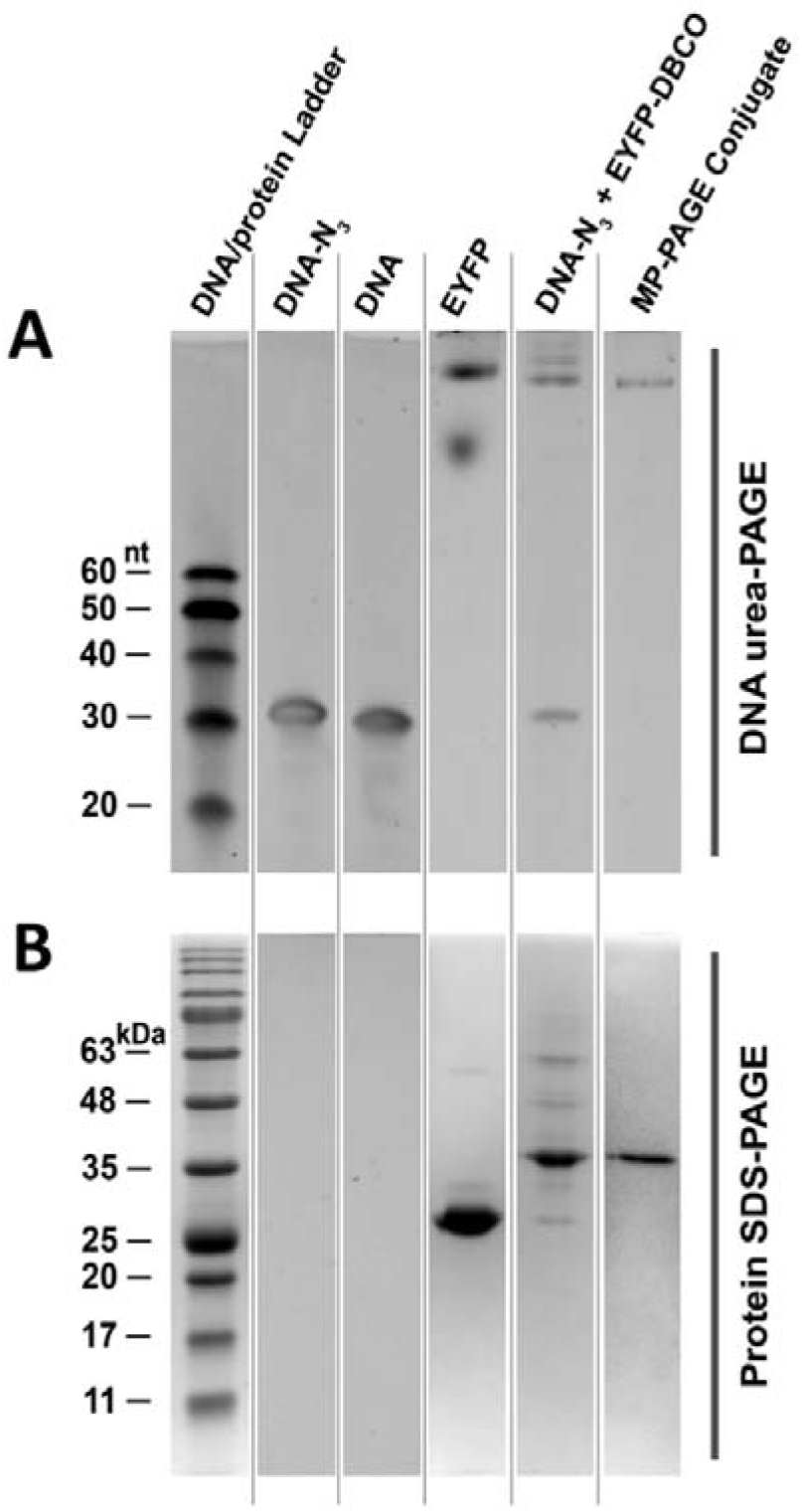
Denaturing PAGE analysis of ssDNA-EYFP conjugate samples. (A) Urea-PAGE profiles showing DNA, EYFP, and conjugate bands. (B) Reducing SDS-PAGE profiles showing EYFP and conjugate bands. DNA-N3 + EYFP-DBCO corresponds to the reaction sample after 4 h incubation followed by filtration using a 30 kDa Amicon centrifugal device.

## Outlook and conclusions

Numerous studies have reported the use of gel electrophoresis for the preparative purification of proteins either in column or slab formats [15,17,20,39,40], These studies relied on either continuous or stopped-flow elution during electrophoresis or methods that require gel slicing after electrophoresis, followed by subsequent extraction steps that limit yield [24]. Although a variety of such apparatuses have become available commercially, they are complex, expensive, or difficult to set up [16]. The MP-PAGE presented herein utilizes standard electrophoretic equipment readily available in most bioanalysis laboratories for purifying different biological samples including proteins, nucleic acids, and bioconjugates. Unlike other preparative gel electrophoresis methods that require high elution volumes and yield diluted protein extracts, MP-PAGE samples are eluted in a small volume (1-2 ml) of buffer that can be collected with a syringe. The MP-PAGE setup also circumvents the need for not only complex gel preparation and shutdown procedures but also specialized equipment such as elution chambers, external pumps, and external fraction collectors.

The versatility of new MP-PAGE method was demonstrated by purifying three distinct samples: a nucleic acid, a protein, and a bioconjugate sample in multicomponent mixtures. The latter sample used in this study could not be purified using existing techniques, particularly due to the low sample amount. Although all the samples used in this work were (or became) fluorescent to allow us to visually track the samples in the gel, the technique is applicable to non-fluorescent samples as well. For example, protein samples could be pre-stained with dyes such as Coomassie G-250 [16] and Instant Band (EZBiolab) and run in parallel to the MP-PAGE to track the position of protein bands and identify the appropriate collection time for the protein.

The MP-PAGE method is convenient for fast and cost-effective, small-volume protein purification in research laboratories. The protein product purity achieved with MP-PAGE is comparable to the purity obtained using chromatography techniques, which are considered as the gold-standard for protein purification. In this study, MP-PAGE purification was performed within 2 h compared to the two-step IMAC+SEC purification that required nearly one day to achieve comparable purities. The ability to tailor the separation resolution by altering the resolving gel concentration further provides a low-cost, tunable approach for purifying proteins of various sizes and charges. In contrast, chromatography techniques require different columns for proteins of different sizes, charges, or metal ion affinities. Compared to IMAC, MP-PAGE also does not require a protein tag for purification, circumventing the need for further protein engineering and overcoming losses from subsequent de-tagging steps. Furthermore, previous studies have shown that native PAGE can be used to separate and purify proteins from intact *E. coli* without disrupting the cells [20]. This ability would allow MP-PAGE to separate proteins in the absence of preliminary cell disruption steps that are necessary for chromatography techniques and can contribute to additional protein loss. We note, however, that these advantages are largely limited to applications that require limited protein amounts. In such cases, chromatography and alternative flow-through systems remain beneficial for larger volume protein purification.

MP-PAGE can also be extended to separate other biomolecules, such as, DNA and DNA-protein conjugates. The MP-PAGE separation demonstrated in this study, for example, was performed under native conditions based on differences in both protein charge and size. MP-PAGE can also be performed in SDS-denaturing conditions, which can separate proteins only based on size. However, though SDS separation benefits from improved separation resolution compared to continuous native-PAGE, this application would be limited to proteins that can be refolded. In addition, this platform can also be applied to specialized biomolecules such as oligonucleotide origami and nanoparticle conjugates. As is the case with proteins and DNA, separation of these conjugates can be achieved by varying concentration, height, and thickness of the resolving gel, as well as buffer composition and applied potential in the electrophoresis device. The use of alternative gel systems, such as discontinues pH gradient or PAG concentration gradient gels, could also be implemented to further tune and optimize separation.

## Supporting Information

Supporting Figs. S1 and S2: chromatograms and SDS-PAGE profiles of SEC and IMAC fractions.

## Acknowledgments

The authors are thankful for the support from the Swiss National Science Foundation Assistant Professor (AP) Energy Grant. They are also thankful for funding support from the Ferdowsi University of Mashhad (Grant No. 3/ 42787).

## References

1. Ersson B, Rydén L, Janson JC. Introduction to Protein Purification. Protein Purification: Principles, High Resolution Methods, and Applications: Third Edition. John Wiley & Sons; 2011. pp. 1–22. doi:10.1002/9780470939932.ch1

2. Cupido T, Pisa R, Kelley ME, Kapoor TM. Designing a chemical inhibitor for the AAA protein spastin using active site mutations. Nat Chem Biol. 2019;15: 444–452. doi:10.1038/s41589-019-0225-6

3. Guyomar C, Thépaut M, Nonin-Lecomte S, Méreau A, Goude R, Gillet R. Reassembling green fluorescent protein for in vitro evaluation of trans-translation. Nucleic Acids Res. 2020;48: E22. doi:10.1093/nar/gkz1204

4. Nickerson JL, Doucette AA. Rapid and Quantitative Protein Precipitation for Proteome Analysis by Mass Spectrometry. J Proteome Res. 2020;19: 2035–2042. doi:10.1021/acs.jproteome.9b00867

5. Foret F, Datinská V, Voráčová I, Novotný J, Gheibi P, Berka J, et al. Macrofluidic Device for Preparative Concentration Based on Epitachophoresis. Anal Chem. 2019;91: 7047–7053. doi:10.1021/acs.analchem.8b05860

6. Goldring JPD. Concentrating proteins by salt, polyethylene glycol, solvent, sds precipitation, three-phase partitioning, dialysis, centrifugation, ultrafiltration, lyophilization, affinity chromatography, immunoprecipitation or increased temperature for protein isolati. Methods in Molecular Biology. Humana Press Inc.; 2019. pp. 41–59. doi:10.1007/978-1-4939-8793-1_4

7. Oberacker P, Stepper P, Bond DM, Höhn S, Focken J, Meyer V, et al. Bio-On-Magnetic-Beads (BOMB): Open platform for high-throughput nucleic acid extraction and manipulation. PLOS Biol. 2019;17: e3000107. Available: https://doi.org/10.1371/journal.pbio.3000107

8. Fekete S, Guillarme D. Ultra-high-performance liquid chromatography for the characterization of therapeutic proteins. TrAC - Trends Anal Chem. 2014;63: 76–84. doi:10.1016/j.trac.2014.05.012

9. Carta G, Jungbauer A. Introduction to Protein Chromatography. Protein Chromatography: Process Development and Scale-Up. John Wiley & Sons; 2020. pp. 63–91. doi:10.1002/9783527824045.ch2

10. Sajjadi SH, Goharshadi EK, Ahmadzadeh H. Heat dissipation in slab gel electrophoresis: The effect of embedded TiO 2 nanoparticles on the thermal profiles. J Chromatogr B Anal Technol Biomed Life Sci. 2019;1118-1119: 63–69. doi:10.1016/j.jchromb.2019.04.030

11. Sajjadi SH, Ahmadzadeh H, Goharshadi EK. Enhanced electrophoretic separation of proteins by tethered SiO2 nanoparticles in an SDS-polyacrylamide gel network. Analyst. 2020;145: 415–423. doi:10.1039/c9an01759c

12. Wu SJ, Schuergers N, Lin KH, Gillen AJ, Corminboeuf C, Boghossian AA. Restriction Enzyme Analysis of Double-Stranded DNA on Pristine Single-Walled Carbon Nanotubes. ACS Appl Mater Interfaces. 2018;10: 37386–37395. doi:10.1021/acsami.8b12287

13. Westermeier R, Gronau S, Becket P, Bülles J, Schickle H, Theßeling G. Electrophoresis in Practice: A Guide to Methods and Applications of DNA and Protein Separations: Fourth Edition. Electrophoresis in Practice: A Guide to Methods and Applications of DNA and Protein Separations: Fourth Edition. John Wiley & Sons; 2005. doi:10.1002/3527603468

14. Sun SM, Hall TC. Application of remazol dye for isolation of protein subunits by preparative SDS-polyacrylamide gel electrophoresis. Anal Biochem. 1974;61: 237–242. doi:10.1016/0003-2697(74)90350-9

15. Naryzhny SN. Upside-down stopped-flow electrofractionation of complex protein mixtures. Anal Biochem. 1996;238: 50–53. doi:10.1006/abio.1996.0249

16. Jian-Hua ZXGC, Lu-Yin Y, Li T, Min W, Dai-Shuang C. Separation and isolation of fusion protein using a new native preparative PAGE device. J Chromatogr Sci. 2012;50: 820–825. doi:10.1093/chromsci/bms077

17. Hao F, Li J, Zhai R, Jiao F, Zhang Y, Qian X. A novel microscale preparative gel electrophoresis system. Analyst. 2016;141: 4953–4960. doi:10.1039/c6an00780e

18. Groschup MH, Boschwitz J, Timoney JF. A convenient gel holder for preparative electrophoretic separation of aggregated bacterial proteins. Electrophoresis. 1991;12: 90–93. doi:10.1002/elps.1150120117

19. Fadouloglou VE. Electroelution of nucleic acids from polyacrylamide gels: A custom-made, agarose-based electroeluter. Anal Biochem. 2013;437: 49–51. doi:10.1016/j.ab.2013.02.021

20. Chew FN, Tan WS, Ling TC, Tey BT. Optimization of a native gel electrophoretic process for the purification of intracellular green fluorescent protein from intact Escherichia coli cells. Process Biochem. 2011;46: 399–403. doi:10.1016/j.procbio.2010.07.032

21. Carpenter HC, Skerritt JH, Wrigley CW, Margolis J. A device for preparative elution electrophoresis using a polyacrylamide gel slab. Electrophoresis. 1986;7: 221–226. doi:10.1002/elps.1150070507

22. Shain DH, Yoo J, Slaughter RG, Hayes SE, Ji TH. Electrofractionation: A technique for detecting and recovering biomolecules. Anal Biochem. 1992;200: 47–51. doi:10.1016/0003-2697(92)90275-C

23. Lim YP, Callanan H, Lin SH, Thompson NL, Hixson DC. Preparative mini-slab gel continuous elution electrophoresis: Application for the separation of two isoforms of rat hepatocyte cell adhesion molecule, cell-CAM 105, and its associated proteins. Anal Biochem. 1993;214: 156–164. doi:10.1006/abio.1993.1471

24. Seelert H, Krause F. Preparative isolation of protein complexes and other bioparticles by elution from polyacrylamide gels. Electrophoresis. 2008;29: 2617–2636. doi:10.1002/elps.200800061

25. Rosen CB, Kodal ALB, Nielsen JS, Schaffert DH, Scavenius C, Okholm AH, et al. Template-directed covalent conjugation of DNA to native antibodies, transferrin and other metal-binding proteins. Nat Chem. 2014;6: 804–809. doi:10.1038/nchem.2003

26. O’Meara TR, O’Meara MJ, Polvi EJ, Pourhaghighi MR, Liston SD, Lin Z-Y, et al. Global proteomic analyses define an environmentally contingent Hsp90 interactome and reveal chaperone-dependent regulation of stress granule proteins and the R2TP complex in a fungal pathogen. PLOS Biol. 2019;17: e3000358. Available: https://doi.org/10.1371/journal.pbio.3000358

27. Zhou Z, Xiang Y, Tong A, Lu Y. Simple and efficient method to purify DNA-protein conjugates and its sensing applications. Anal Chem. 2014;86: 3869–3875. doi:10.1021/ac4040554

28. Yang L, Bui L, Hanjaya-Putra D, Bruening ML. Membrane-Based Affinity Purification to Identify Target Proteins of a Small-Molecule Drug. Anal Chem. 2020;92: 11912–11920. doi:10.1021/acs.analchem.0c02316

29. Zubkovs V, Wu SJ, Rahnamaee SY, Schuergers N, Boghossian AA. Sitespecific protein conjugation onto fluorescent single-walled carbon nanotubes. Chem Mater. 2020;32: 8798–8807. doi:10.1021/acs.chemmater.0c02051

30. Maxam AM, Gilbert W. A new method for sequencing DNA. Proc Natl Acad Sci U S A. 1977;74: 560–564. doi:10.1073/pnas.74.2.560

31. De Meulenaere E, Nguyen Bich N, De Wergifosse M, Van Hecke K, Van Meervelt L, Vanderleyden J, et al. Improving the second-order nonlinear optical response of fluorescent proteins: The symmetry argument. J Am Chem Soc. 2013;135: 4061–4069. doi:10.1021/ja400098b

32. Green MR, Sambrook J. Isolation of DNA fragments from polyacrylamide gels by the crush and soak method. Cold Spring Harb Protoc. 2019;2019: 143–146. doi:10.1101/pdb.prot100479

33. Snapp EL, McCaul N, Quandte M, Cabartova Z, Bontjer I, Källgren C, et al. Structure and topology around the cleavage site regulate post-translational cleavage of the HIV-1 gp160 signal peptide. Gilmore R, editor. Elife. 2017;6: e26067. doi:10.7554/eLife.26067

34. Saccà B, Niemeyer CM. Functionalization of DNA nanostructures with proteins. Chem Soc Rev. 2011;40: 5910–5921. doi:10.1039/c1cs15212b

35. Singh Y, Murat P, Defrancq E. Recent developments in oligonucleotide conjugation. Chem Soc Rev. 2010;39: 2054–2070. doi:10.1039/b911431a

36. Yang YR, Liu Y, Yan H. DNA Nanostructures as Programmable Biomolecular Scaffolds. Bioconjug Chem. 2015;26: 1381–1395. doi:10.1021/acs.bioconjchem.5b00194

37. Khatwani SL, Kang JS, Mullen DG, Hast MA, Beese LS, Distefano MD, et al. Covalent protein-oligonucleotide conjugates by copper-free click reaction. Bioorganic Med Chem. 2012;20: 4532–4539. doi:10.1016/j.bmc.2012.05.017

38. Lapiene V, Kukolka F, Kiko K, Arndt A, Niemeyer CM. Conjugation of fluorescent proteins with DNA oligonucleotides. Bioconjug Chem. 2010;21: 921–927. doi:10.1021/bc900471q

39. Laremore TN, Ly M, Solakyildirim K, Zagorevski D V., Linhardt RJ. High-resolution preparative separation of glycosaminoglycan oligosaccharides by polyacrylamide gel electrophoresis. Anal Biochem. 2010;401: 236–241. doi:10.1016/j.ab.2010.03.004

40. Gabe CM, Brookes SJ, Kirkham J. Preparative SDS PAGE as an alternative to his-tag purification of recombinant Amelogenin. Front Physiol. 2017;8: 424. doi:10.3389/fphys.2017.00424

