## Supplemental Figs. 1 and 2 for "A simple micropreparative gel electrophoresis technique for purification of proteins, nucleic acids, and bioconjugates"

^a^ École Polytechnique Fédérale de Lausanne (EPFL), CH-1015, Lausanne, Switzerland

^b^ Chemistry Department, Faculty of Science, Ferdowsi University of Mashhad, Mashhad 9177948974, Iran

*

*

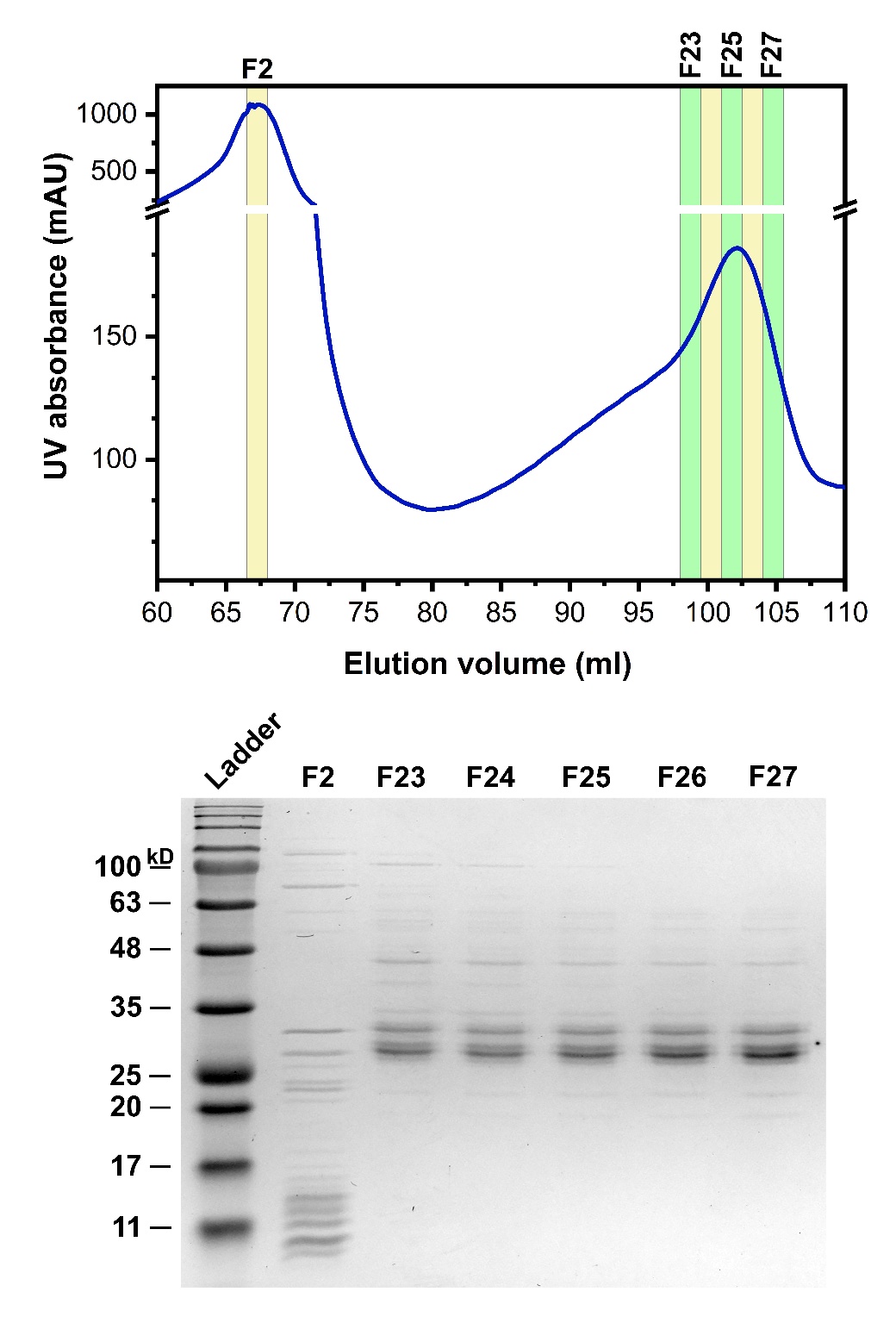

**Fig S1**. Chromatogram (top) and Coomassie Brilliant Blue (CB)-stained reducing SDS-PAGE profiles (bottom) of fractions eluted from the SEC column.

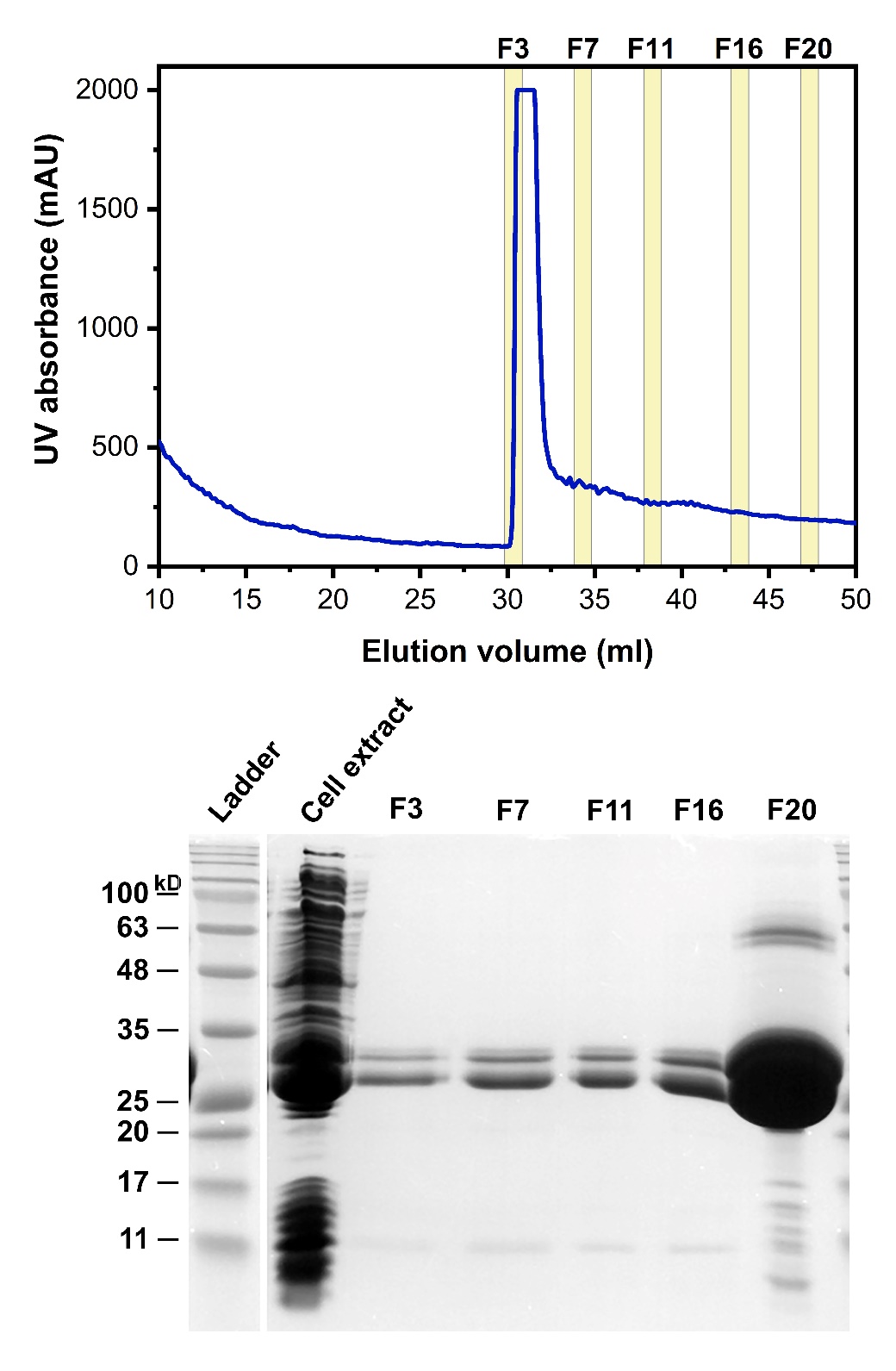

**Fig S2**. Chromatogram (top) and Coomassie Brilliant Blue (CB)-stained reducing SDS-PAGE profiles (bottom) of fractions eluted from the IMAC column.
